# Gasdermin E couples mitochondrial damage to pyroptotic neurodegeneration in Parkinson’s disease

**DOI:** 10.64898/2025.12.14.694264

**Authors:** Jiasong Pan, Lu Geng, Xuehe Liu, Yinglei Wang, Ning Fang, Feiyan Xie, Yajie Shen, Yifan Deng, Yuyun Han, Suhua Li, Wenyong Xiong, Hua He, Ningyi Shao, Jian Wang, Jixi Li

**Affiliations:** Department of Neurology, Huashan Hospital and School of Life Sciences, State Key Laboratory of Genetics and Development of Complex Phenotypes, Shanghai Engineering Research Center of Industrial Microorganisms, Fudan University, Shanghai, 200438, China; Wisdom Lake Academy of Pharmacy, Xi’an Jiaotong-Liverpool University, Jiangsu, 215123, China; Division of Natural Science, Duke Kunshan University, Jiangsu, 215316, China; Key Laboratory of Medicinal Chemistry for Natural Resource, Ministry of Education; Yunnan Provincial Center for Research & Development of Natural Products; School of Pharmacy, Yunnan University, Kunming, 650500, China; Department of Neurosurgery, The Third Affiliated Hospital, Naval Medical University, Shanghai, 200433, China; Department of Biomedical Sciences, Faculty of Health Sciences, University of Macau, Macau, China; Department of Neurology and National Center for Neurological Diseases, Huashan Hospital, Fudan University, Shanghai, 200040, China

**Author notes:** J.P. and L.G. contributed equally to this work.

**Keywords:** Parkinson’s disease, GSDME, Mitochondrial dysfunction, Pyroptosis

## Abstract

Parkinson’s disease (PD) features progressive loss of nigrostriatal dopamine neurons, but how mitochondrial damage engages programmed cell death pathways remains unresolved. Here, we identify gasdermin E (GSDME), the caspase-3-activated executor of pyroptosis, as a critical mediator of neurodegeneration in toxin-based PD models. In primary neurons and SH-SY5Y cells, the mitochondrial complex I inhibitor MPTP/MPP⁺ triggered caspase-3 activation, GSDME cleavage, and lytic membrane rupture. Genetic silencing of Gsdme or its transcriptional regulator SP1 reduced neuronal death. *In vivo*, Gsdme deficiency preserved substantia nigra pars compacta dopaminergic neurons, improved motor performance, and mitigated anxiety- and depression-like behaviors after MPTP administration. Loss of Gsdme also dampened microglial and astrocytic activation and lowered proinflammatory cytokines in striatum and substantia nigra. Mechanistically, cleaved GSDME localized to mitochondria, disrupted membrane potential, increased reactive oxygen species, and precipitated organelle injury, thereby coupling mitochondrial dysfunction to pyroptotic cell death. These findings identify GSDME-mediated pyroptosis as a mechanistic link between mitochondrial toxicity and neuroinflammation in PD and nominate GSDME as a therapeutic entry point to slow disease progression.

**Significance:** How mitochondrial injury kills dopamine neurons in Parkinson’s disease is a central unresolved question. We show that the pyroptosis executor GSDME is required for neurodegeneration and neuroinflammation in MPTP models, mechanistically linking caspase-3 activation and mitochondrial damage to lytic cell death. Targeting GSDME may provide a strategy to protect vulnerable neurons in PD.

**Graphic Abstract:** 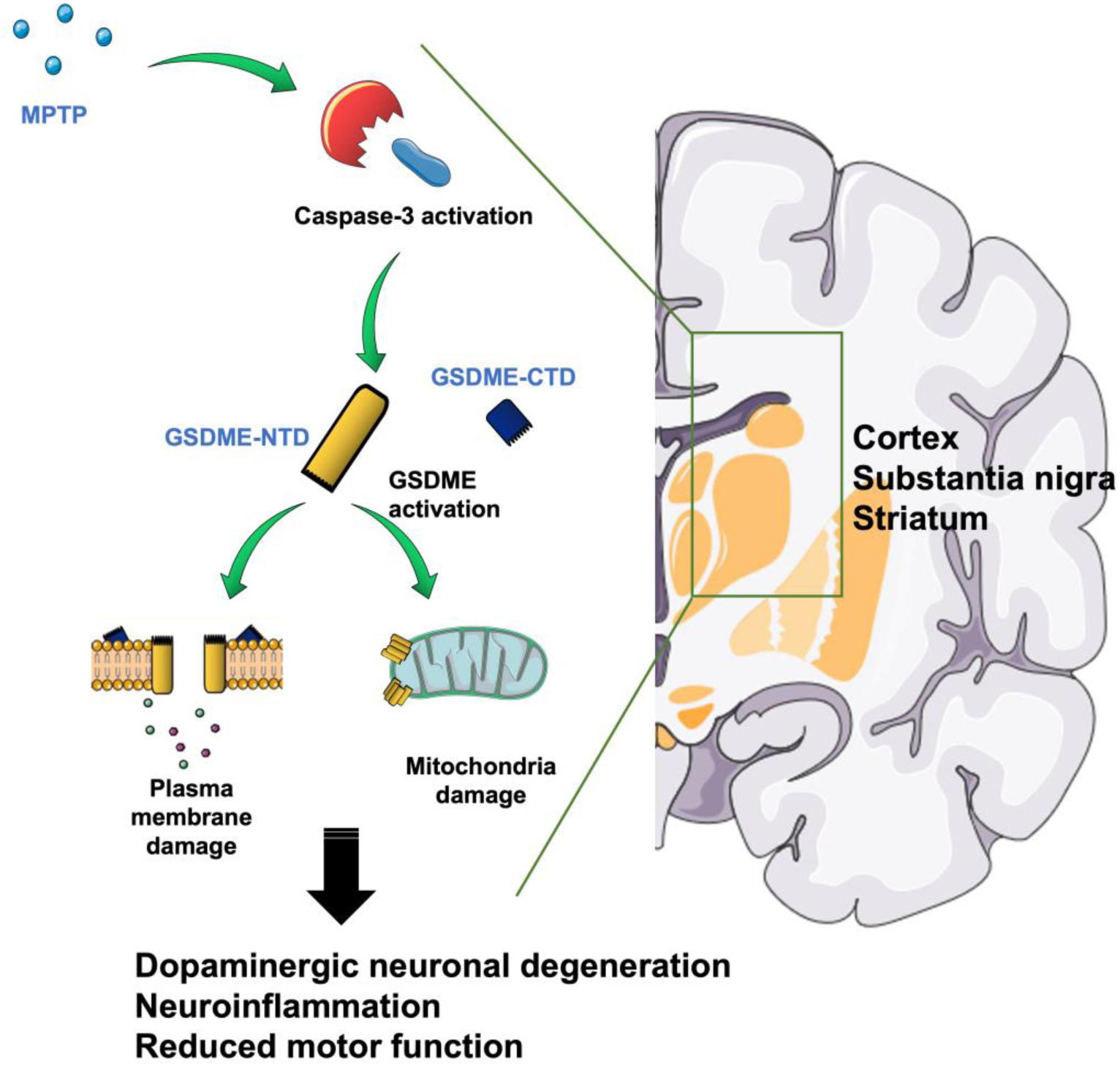

**Highlights:** GSDME drives PD-linked neuronal loss. Silencing GSDME (or its transcriptional factor Sp1) ameliorates MPTP-triggered neuronal cell death.

Genetic ablation of Gsdme is protective in the MPTP-induced PD mouse model. *Gsdme*^-/-^mice retain nigrostriatal DA neurons, show improved motor and affective behaviors, and exhibit reduced microglial and astrocytic activation and cytokines release upon MPTP treatment.

Cleaved GSDME accumulates on mitochondria, collapses ΔΨm, raises ROS, and links mitochondrial toxicity to pyroptosis, thus positioning GSDME as a tractable target to slow PD progression.

## Introduction

Parkinson’s disease (PD) is a progressive neurodegenerative disorder that has become a growing public health challenge in aging populations ^1^. Clinically, PD is defined by its cardinal motor features, including bradykinesia, resting tremor, rigidity, and postural instability, as well as a range of non-motor disturbances including hyposmia, REM sleep behavior disorder, and cognitive decline ^2,3^. Pathologically, PD is marked by the loss of dopaminergic neurons in the substantia nigra pars compacta (SNpc), leading to striatal dopamine depletion ^4^. Mutations in *SNCA*, which encodes α-synuclein, are a major genetic cause of Parkinson’s disease. Misfolded α-synuclein assembles into fibrils and Lewy bodies that drive cellular toxicity and neuronal death ^5,6^. Environmental toxicants including rotenone, organochlorine pesticides, 6-hydroxydopamine (6-OHDA), MPTP, and carbon tetrachloride can also precipitate dopaminergic degeneration through mitochondrial disruption and neuroinflammation, independent of Lewy body pathology ^7,8^. Although pharmacological and surgical therapies alleviate symptoms, no treatment halts or reverses disease progression, underscoring the need to define the cellular and molecular mechanisms that couple mitochondrial injury to neuron loss.

Programmed neuronal death contributes to neurodegeneration through apoptosis, necroptosis, and pyroptosis. In Parkinson’s disease, mitochondrial damage is a well-established trigger of apoptotic pathways ^9,10^. Necroptosis mediated by RIPK1, RIPK3, and MLKL has recently been implicated in disease progression, and Mlkl ablation ameliorates motor deficits in SNCA A53T mouse models ^11^. Pyroptosis mediated by the gasdermin family has emerged as another relevant mechanism ^12^. Gasdermin E (GSDME) is cleaved by caspase-3 to release an N-terminal fragment that oligomerizes in membranes and causes lytic rupture ^13,14^. First recognized as an antitumor effector that redirects apoptosis to pyroptosis ^14,15^, GSDME is now linked to several neurological disorders. GSDME is upregulated in neurons of APP/PS1 models of Alzheimer’s disease where it promotes Aβ-induced toxicity and cognitive impairment ^16^. In amyotrophic lateral sclerosis (ALS), GSDME mediates mitochondrial damage, axonal degeneration, and disease progression upon caspase-3 activation by pathogenic protein aggregates, and its genetic ablation prolongs survival and preserves motor function ^17^. Furthermore, GSDME majorly expresses in neurons, whereas its family protein GSDMD expresses in microglia ^17^. Also, GSDME has been linked to HIV-associated neurological complications ^18^. Despite these observations, whether GSDME contributes to Parkinson’s disease remains unknown.

Here, we demonstrate that GSDME is a critical executor of dopaminergic neurodegeneration in PD. Neurotoxins 6-OHDA and MPTP activated GSDME-dependent pyroptosis and mitochondrial damage in primary neurons, an effect mitigated by depletion Gsdme or its transcriptional factor Sp1. In MPTP-induced mice, Gsdme deficiency improves motor performance, reduces anxiety- and depression-like behaviors, preserves nigrostriatal dopaminergic neurons, and dampens microglial and astrocytic activation. RNA sequencing reveals that Gsdme loss reverses mitochondrial stress and innate immune signatures. These findings establish GSDME-driven pyroptosis as a mechanistic link between mitochondrial dysfunction, neuroinflammation, and neuronal loss in Parkinson’s disease and nominate GSDME as a therapeutic entry point.

## Results

### PD-relevant stressors activate GSDME and drive neuronal death

To test whether Parkinsonian insults engage GSDME, we exposed primary mouse cortical neurons and human SH-SY5Y cells to MPTP or 6-OHDA ^19,20^. Both toxins markedly increased GSDME cleavage, yielding the pore-forming N-terminal fragment (GSDME-NT) and inducing lytic death (Fig. 1A-E and S1). Silencing Gsdme or its transcriptional regulator SP1 ^21^ significantly reduced toxin-induced death in both cell types and diminished GSDME-NT generation (Fig. 1A-E). Pharmacological blockade of caspase-3 activity with Z-DEVD-FMK suppressed GSDME cleavage in a dose-dependent manner and mitigated neurotoxin-induced cell death in primary neurons and SH-SY5Y cells (Fig. 1C-D and S1), indicating that caspase-3-dependent GSDME activation is a central mechanism of pyroptosis in this context ^21,22^.

**Figure 1.**
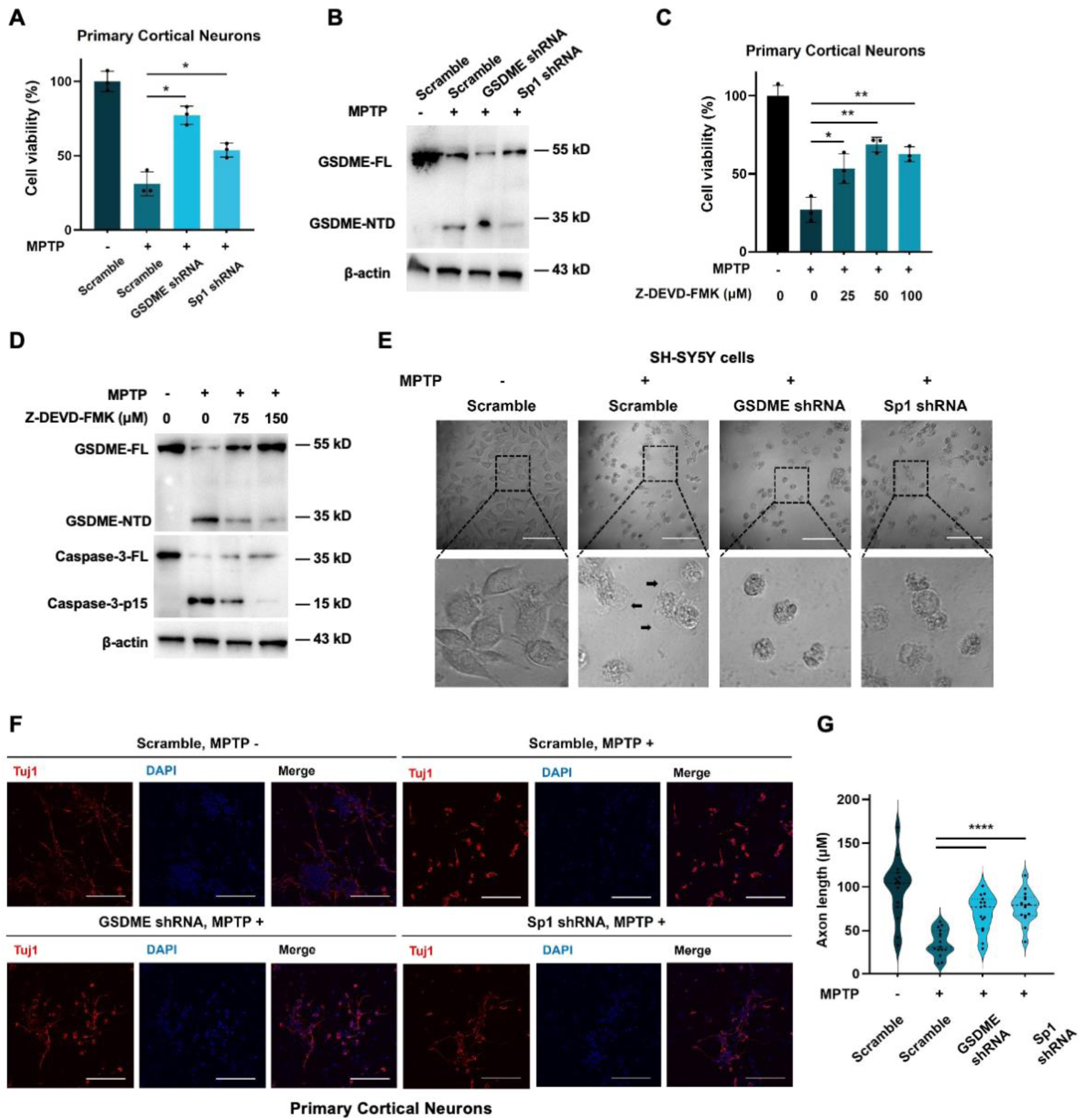
Genetic or pharmacological inhibition of GSDME limits neurotoxin-triggered pyroptosis and axonal injury. (**A-B**) Primary mouse cortical neurons were transduced with lentivirus encoding scrambled control, GSDME shRNA, or Sp1 shRNA. After 72 h, cultures were challenged with MPTP as indicated. Cell viability was quantified 24 h later (normalized to vehicle) (**A**). Immunoblots were performed for full-length GSDME (GSDME-FL), cleaved N-terminal GSDME (GSDME-NTD), and β-actin (**B**). (**C-D**) Neurotoxin-treated primary cortical neurons were exposed to increasing concentrations of the caspase-3 inhibitor Z-DEVD-FMK, and cell viability was quantified (**C**). Immunoblot analyses of GSDME-FL, GSDME-NTD, caspase-3-FL, and caspase-3-p15 (with β-actin as a loading control) in Z-DEVD-FMK-treated, neurotoxin-stimulated cells (**D**). (**E**) Representative images of cell morphology following MPTP exposure in SH-SY5Y cultures expressing scrambled, GSDME-, or Sp1-shRNAs. Black arrows showing the blebbing. Scale bar, 100 µm. (**F-G**) Primary cortical neurons stained for neuronal marker Tuj1 (red) after MPTP treatment. Axonal length was quantified blind to condition. Scale bar, 100 µm. All experiments were performed in biological triplicates. Data represent the mean ± SD. ns, not significant; *P < 0.05; **P < 0.01; ***P < 0.001; ****P < 0.0001.

Morphologically, MPTP-treated cells exhibited characteristic pyroptotic features, including cellular swelling and large plasma membrane blebs, whereas GSDME- or Sp1-depleted cells instead displayed apoptotic condensation and fragmented nuclei (Fig. 1E). In primary cortical neurons, MPTP elicited marked axonal retraction and loss of synaptic structure, mimicking the neurodegenerative changes observed in PD patients ^23^, whereas GSDME or Sp1 silencing preserved axonal integrity and prevented synapse loss (Fig. 1F-G). Together, these findings identify GSDME-mediated pyroptosis as a pivotal contributor to neuronal damage induced by PD-relevant toxins.

### GSDME ablation alleviates motor and behavioral deficits in MPTP-induced mice

To investigate whether GSDME functions in PD pathogenesis, we used the acute MPTP model, in which systemic MPTP inhibits mitochondrial complex I, increases ROS, and causes nigrostriatal neuron loss and motor/neuropsychiatric impairments ^20^. Wild-type (WT) and *Gsdme*^−/−^ littermates received MPTP or saline and were assessed behaviorally (Fig. 2A). In the tail-suspension test, *Gsdme*^−/−^ mice exhibited reduced immobility after MPTP treatment compared with WT, whereas saline-treated groups were indistinguishable (Fig. 2B). Open-field analysis showed comparable baseline locomotion and center exploration between genotypes, but MPTP significantly decreased total distance and center entries in WT mice. Both measures were preserved in *Gsdme*^−/−^ mice (Fig. 2C-E). Consistently, Gsdme deficiency improved motor coordination and bradykinesia endpoints: rotarod latency was longer and pole-test descent times were shorter in *Gsdme*^−/−^ mice relative to WT after MPTP treatment (Fig. 2F-G).

**Figure 2.**
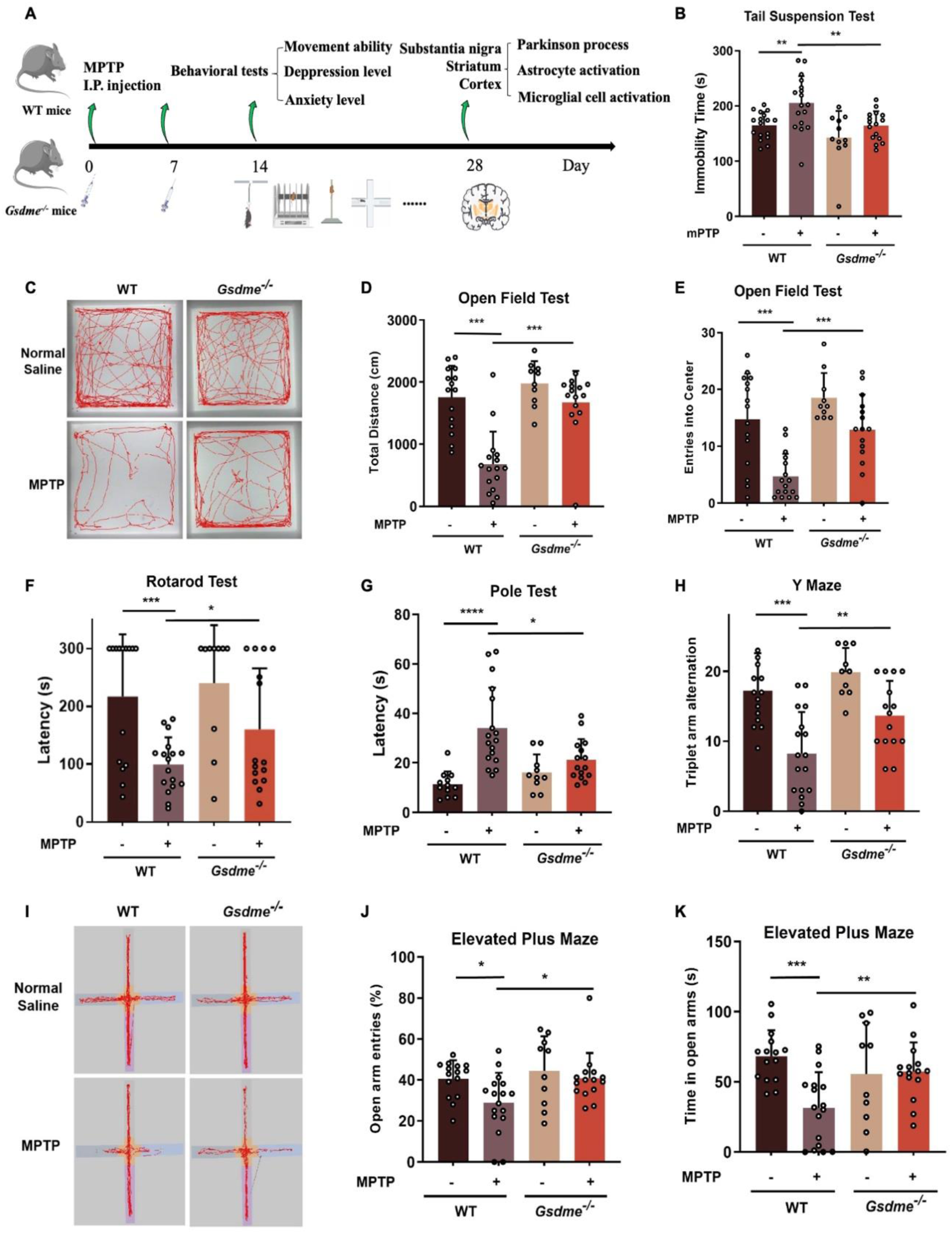
Gsdme deficiency ameliorates motor deficits, anxiety- and depression-like behaviors in MPTP-treated mice. (**A**) Schematic workflow of the experimental design, including MPTP injections and subsequent behavioral and histopathological assessments. (**B**) Total immobility time in the tail suspension test. (**C-E**) Open-field test: representative locomotor traces (**C**), total distance traveled (**D**), and total entries into the central area (**E**) for WT-MPTP (–), WT-MPTP (+), *Gsdme*^−/−^-MPTP (–), and *Gsdme*^−/−^-MPTP (+) mice. (**F**) Latency to fall in the accelerating rotarod test. (**G**) Time required to descend a vertical pole. (**H-I**) Elevated plus-maze test: representative movement traces, quantification of entries, and time spent in the open arms. (**J**) Number of spontaneous alternations among the three arms of the Y-maze test. All experiments were performed in biological triplicates. Data represent the mean ± SD. ns, not significant; *P < 0.05; **P < 0.01; ***P < 0.001; ****P < 0.0001.

To probe affective and cognitive domains, we used the elevated plus maze and Y-maze methods. Following MPTP treatment, *Gsdme*^−/−^ mice spent more time and made more entries into open arms, and displayed higher spontaneous alternation rates than WT controls, indicating reduced anxiety-like behavior and improved working memory (Fig. 2H-I). Together, these data show that loss of Gsdme mitigates both motor and neuropsychiatric deficits in the MPTP model, supporting a pathogenic role for GSDME *in vivo*.

### Gsdme deficiency preserves nigrostriatal neurons and dampens neuroinflammation

Next, we examined neuropathology in substantia nigra (SN), striatum, and cortex after acute MPTP. In WT mice, tyrosine hydroxylase (TH) staining showed marked loss of SNpc dopaminergic neurons and depletion of TH⁺ striatal projections. *Gsdme*^−/−^ littermates exhibited partial preservation of both neuronal populations (Fig. 3A-B). Immunoblot analyses further confirmed robust caspase-3 and GSDME cleavage in the SN, striatum, and cortex of MPTP-treated WT mice (Fig. 3C). Notably, caspase-3 cleavage remained unchanged in *Gsdme*^−/−^ mice, although TH expression was significantly preserved and α-synuclein phosphorylation was reduced compared with WT controls (Fig. 3C).

**Figure 3.**
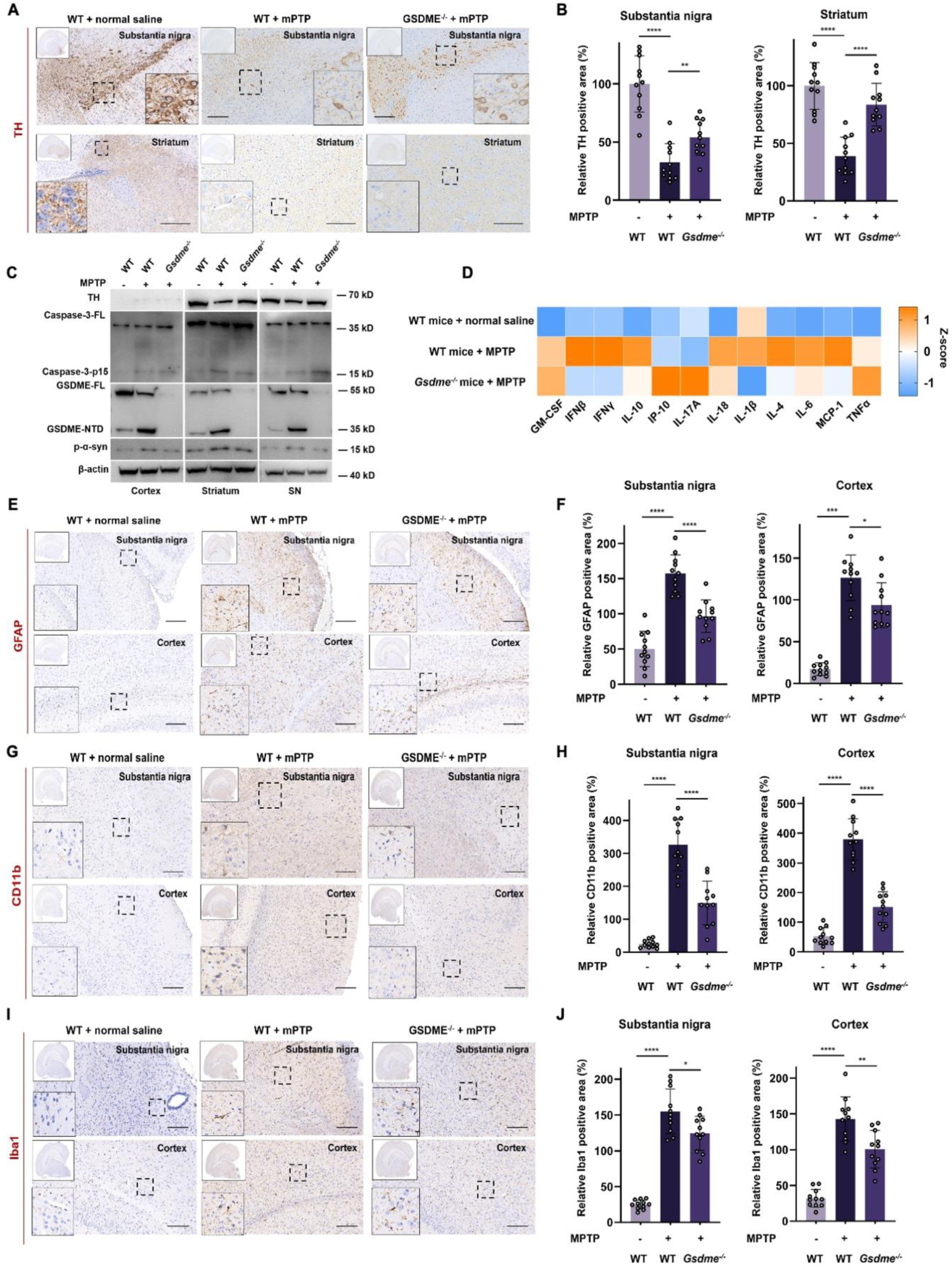
GSDME ablation preserves nigrostriatal dopaminergic neurons and dampens neuroinflammation. (**A-B**) Tyrosine hydroxylase (TH) immunostaining in substantia nigra (SN) and striatum from WT-MPTP (−), WT-MPTP (+), and *Gsdme*^−/−^-MPTP (+) mice. (**A**) Whole-section overviews (top left) with higher-magnification views of boxed regions below. Scale bars, 200 µm. (**B)** Quantification of TH⁺ area in striatum and SN. (**C**) Immunoblot analyses of TH, GSDME-FL, GSDME-NTD, caspase-3-FL, caspase-3-p15, p-α-synuclein, and β-actin in SN, striatum, and cortex from the indicated groups. (**D-I**) Representative immunostaining for astrocytes (GFAP) and microglia (CD11b, Iba1) in SN and cortex (overview at top left; higher magnification of boxed regions at bottom left). Scale bars, 200 µm. (**E, G, I**) Quantification of GFAP-, CD11b-, and Iba1-positive area. (**J**) Heat map of Z-scored serum cytokine profiles in the indicated groups, with orange representing increased and blue representing decreased concentrations.

Furthermore, we assessed glial activation and neuroinflammation. ELISA analyses revealed significant elevations in the serum concentrations of IFN-β, IFN-γ, IL-10, IL-18, IL-1β, IL-4, IL-6, and MCP-1 following MPTP treatment in WT mice, which were effectively suppressed in *Gsdme*^−/−^ littermates (Fig. 3D). In MPTP-treated WT mice, GFAP immunoreactivity was elevated in the SN and cortex, indicative of astrogliosis, whereas *Gsdme*^−/−^ mice showed attenuated GFAP expression (Fig. 3E-F). Similarly, microglial activation markers CD11b and Iba1 were upregulated in the SN and cortex of WT PD mice upon MPTP treatment, but this response was blunted in *Gsdme*^−/−^ mice (Fig. 3G-J). Together, these results demonstrate that GSDME deficiency preserves nigrostriatal dopamine circuitry and alleviates neuroinflammation, highlighting its pivotal role in PD pathogenesis.

### RNA-seq analysis reveals Gsdme loss reverses mitochondrial stress and innate immune signatures

To define how GSDME deficiency reshapes the PD process, we performed RNA-seq on substantia nigra from control wild-type (WT), MPTP-exposed WT (WT-MPTP), and MPTP-exposed *Gsdme*^−/−^ (*Gsdme*^−/−^-MPTP) mice. Comparing WT-MPTP with WT revealed a robust transcriptional response (DESeq2, FDR < 0.05): 135 genes were upregulated and 114 were downregulated (Fig. 4A). Among the most induced transcripts were cortical or forebrain developmental regulators (Ddn, Foxg1, Dlx6, and Dlx6os1), suggesting maladaptive reactivation of developmental programs after toxin injury. Conversely, genes involved in lipid homeostasis, stress responses, and cell-cycle control (Plin4, Prl, Cdkn1a, and Gm45579) were repressed. Gene Ontology (GO) enrichment highlighted neuronal development and synaptic plasticity terms, whereas KEGG analysis pointed to cholinergic synapse, neuroactive ligand–receptor interaction, and calcium signaling-pathways linked to excitotoxicity and neuroinflammation in PD (Fig. 4C and 4E).

**Figure 4.**
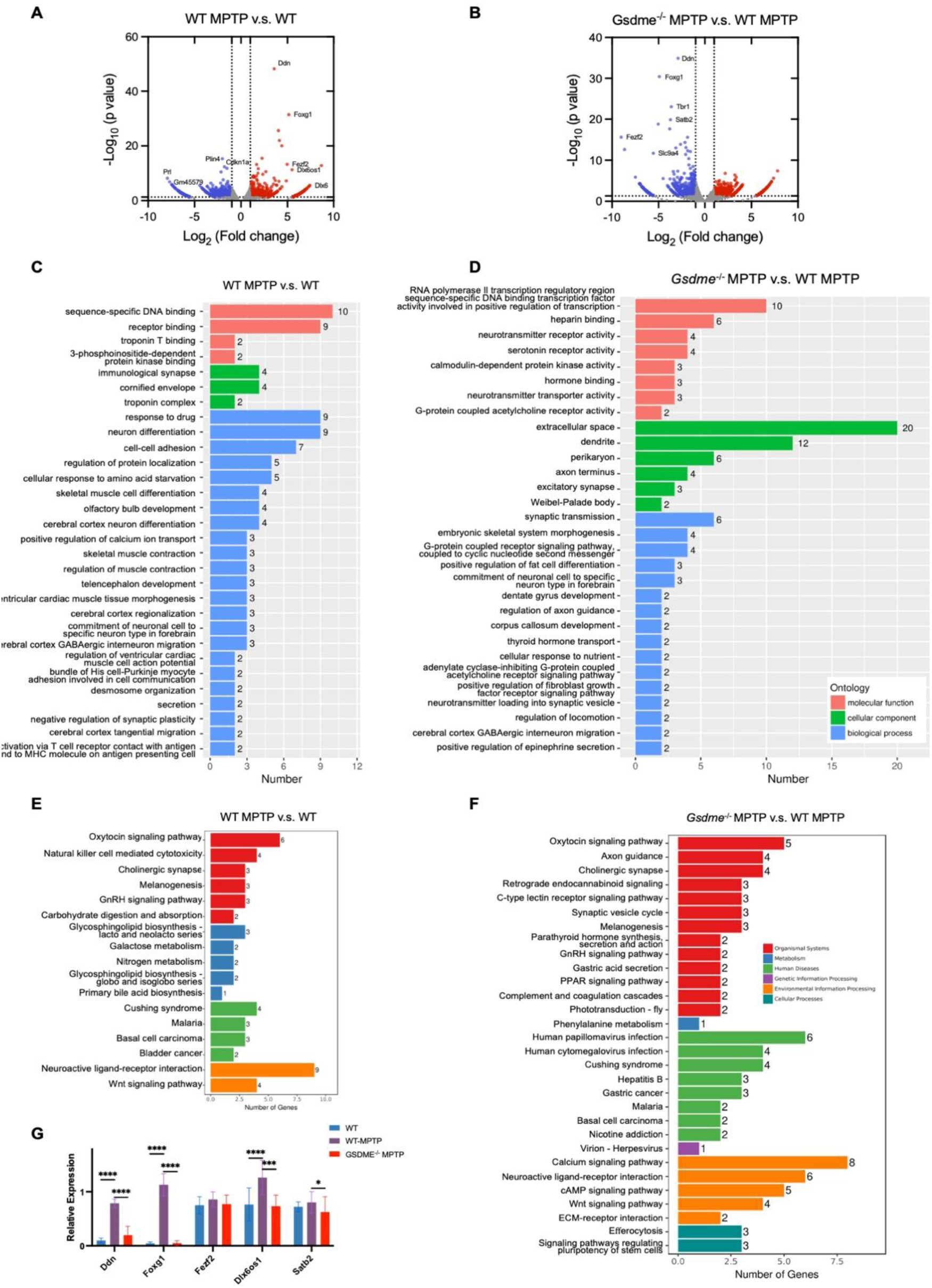
GSDME deficiency reverses the MPTP-induced pathogenic transcriptional program in substantia nigra. **(A-B)** Volcano plots showing differentially expressed genes (DEGs) in the substantia nigra. WT-MPTP versus WT (**A**); *Gsdme*^−/−^-MPTP versus WT-MPTP **(B)**. Red/blue dots denote significantly up-/down-regulated genes, respectively (∣log2FC∣>1, FDR<0.05). **(C-D)** Top 20 enriched Gene Ontology (GO) Biological Process terms for the identified DEGs in WT-MPTP vs. WT **(C)** and in *Gsdme*^−/−^-MPTP vs. WT-MPTP **(D)**, ranked by P-value. (**E-F)** Top enriched KEGG pathways in WT-MPTP vs. WT **(E)** and in *Gsdme*^−/−^-MPTP vs. WT-MPTP **(F)**. **(G)** Validation of key differentially expressed genes by quantitative real-time PCR (qPCR). Data are normalized to a housekeeping gene (GAPDH) and presented as mean ± SEM. (n=4 per group; ∗P<0.05, ∗∗P<0.01, ∗∗∗P<0.001).

Strikingly, in *Gsdme*^−/−^-MPTP versus WT-MPTP, 157 genes were differentially expressed (45 up, 112 down; FDR < 0.05), with broad reversal of the developmental signature: Foxg1, Fezf2, Tbr1, Satb2, Ddn, and Dlx6os1 were significantly reduced (Fig. 4B). GO terms related to GABAergic interneuron migration, axon guidance, and dentate gyrus development were dampened (Fig. 4D), together with immune or glial programs (for example, positive regulation of FGF receptor signaling and neuroinflammatory response). KEGG analysis showed attenuated Wnt signaling, ECM-receptor interaction, and retrograde endocannabinoid signaling (Fig. 4F), consistent with reduced gliosis, synaptic destabilization and neuroimmune crosstalk. Next, represented transcripts (Foxg1, Fezf2, Satb2, and Crym) were validated using the qPCR method (Fig. 4G). Together, MPTP elicits a broad, potentially maladaptive reprogramming of developmental and neuroimmune networks in the midbrain. Gsdme deficiency blunts these responses and normalizes metabolic and synaptic pathways, indicating that beyond executing pyroptosis, GSDME shapes transcriptional states that couple mitochondrial injury to neuroinflammation and circuit remodeling in PD.

### Activated GSDME localizes to mitochondria and amplifies mitochondrial injury

MPTP exerts toxicity by entering mitochondria, inhibiting complex I, lowering ATP, and elevating reactive oxygen species (ROS) ^20^. In SH-SY5Y cells, MPTP converted the diffuse cytosolic pattern of GSDME to a punctate signal that colocalized with the mitochondrial marker TOM20, indicating recruitment of activated GSDME to mitochondria (Fig. 5A). In primary cortical neurons and SH-SY5Y cells, MPTP caused loss of mitochondrial membrane potential (TMRE) and increased ROS; both effects were significantly attenuated by GSDME or SP1 knockdown (Fig. 5B-E). Ultrastructurally, GSDME depletion preserved mitochondrial length and cristae and reduced fragmentation following MPTP treatment observed by transmission electron microscopy (Fig. 5F-G). Consistently, GSDME knockdown or caspase-3 inhibition with Z-DEVD-FMK prevented MPTP-induced loss of TOM20 and upregulation of Fis1, indicative of improved mitochondrial dynamics and integrity ^24^ (Fig. 5H).

**Figure 5.**
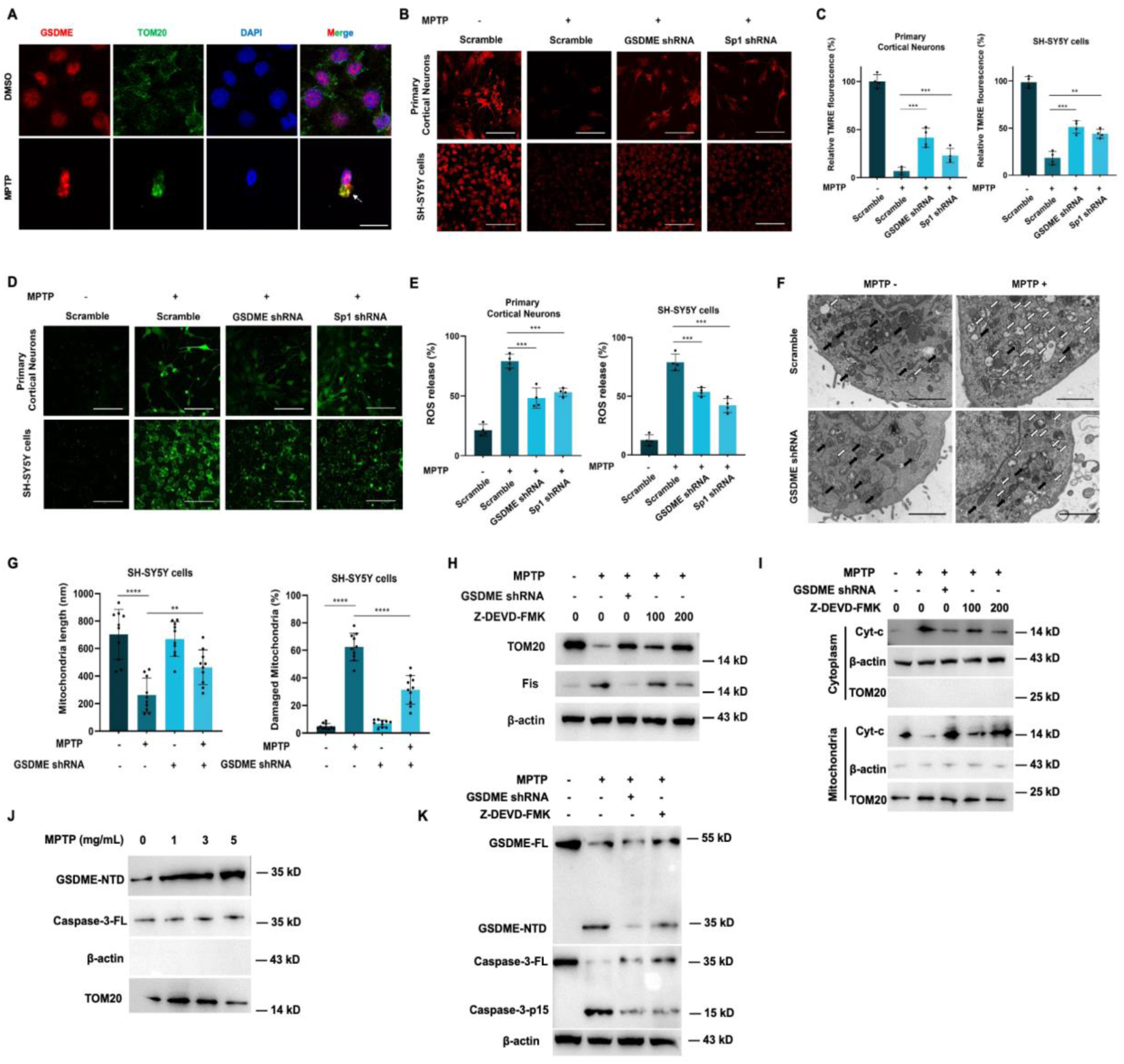
GSDME drives mitochondrial damage and ROS in MPTP-treated neurons. **(A)** Confocal co-localization of GSDME (red) with the mitochondrial marker TOM20 (green) in SH-SY5Y cells treated with or without MPTP (Scale bar: 20 μm). (**B-C)** Mitochondrial membrane potential assessed by TMRE in SH-SY5Y cells and primary cortical neurons transduced with scrambled, GSDME, or Sp1 shRNAs and exposed to MPTP; representative images (**B**, scale bar, 100 µm) and quantification normalized to DMSO (**C**). (**D-E)** Cellular ROS measured by DCFH-DA fluorescent probes staining for scramble/GSDME/SP1-shRNA transfected SH-SY5Y cell and mice primary cortical neurons stimulated by MPTP, scale bars, 100 μm (**D**). The quantification results of ROS release normalized to DMSO controls are shown in **E**. (**F-G)** Transmission electron microscopy of SH-SY5Y cells (scrambled vs GSDME shRNA) treated with either DMSO or MPTP showing intact (black arrows) and damaged (white arrows) mitochondria (scale bar, 500 nm) (**F**) with quantification of mitochondrial length and damage ratio (**G**). (**H**) Immunoblot of TOM20, Fis1, and β-actin in SH-SY5Y cells with GSDME knockdown or caspase-3 inhibition (Z-DEVD-FMK) stimulated by MPTP. (**I**) SH-SY5Y cells were transfected with GSDME shRNA or treated by Z-DEVD-FMK, followed by MPTP stimulation. Mitochondria were isolated and analyzed together with the remaining cellular fractions by Western blot to detect the levels of Cyt-C, β-actin and TOM20. (**J**) Detection of mitochondrial GSDME-NTD and caspase-3 after MPTP treatment. TOM20 and β-actin are as markers. (**K**) Western blotting analysis of protein levels (GSDME-FL, GSDME-NTD, Caspase-3-FL, Caspase-3-p15, Cyt-C and β-actin) in GSDME shRNA transfected or Z-DEVD-FMK treated SH-SY5Y cells stimulated by MPTP. All experiments were performed in biological triplicates. Data represent the mean ± SD. ns, not significant; *P < 0.05; **P < 0.01; ***P < 0.001; ****P < 0.0001.

Mitochondrial damage promotes cytochrome-c (Cyt-c) release ^25^, caspase-3 activation, and GSDME cleavage, suggesting a feed-forward loop. Immunoblot analyses confirmed that MPTP increased Cyt-C release, an effect significantly blunted by GSDME silencing or caspase-3 inhibition (Fig. 5I). Notably, GSDME ablation also suppressed caspase-3 activation, suggesting that GSDME promotes upstream caspase-3 activity. Next, mitochondrial fractionation assays were performed to define the subcellular localization of activated GSDME. Upon MPTP exposure, cleaved GSDME-NTD accumulated within the mitochondrial compartment and colocalized with the mitochondrial marker TOM20, whereas caspase-3 was minimally associated with mitochondria (Fig. 5J-K). Together, these findings reveal that cleaved GSDME-NTD translocates to mitochondria to amplify mitochondrial damage and drive a feed-forward caspase-3-GSDME-Cyt-C loop that promotes neuronal death.

### Cleaved GSDME is required for mitochondrial injury and neuronal pyroptosis in the MPTP model

To test the necessity of GSDME cleavage, we reconstitute either wild-type (WT) GSDME or a cleavage-resistant GSDME-D270A mutant in primary cortical neurons obtained from *Gsdme*^−/−^ mice. The D270A point mutation ablates the caspase-3 cleavage site, yielding a constitutively inactive GSDME ^12^. Upon MPTP treatment, GSDME-WT reconstitution restored vulnerability to mitochondrial and cellular injury, evidenced by increased LDH release and reduced viability, whereas GSDME-D270A had no significant effect (Fig. 6A-B). Consistently, GSDME-WT, but not the D270A variant, reinstated the loss of mitochondrial membrane potential and elevated ROS generation following MPTP exposure (Fig. 6C-F). Immunoblot analyses confirmed that GSDME-WT reconstitution restored GSDME-NTD generation and enhanced caspase-3 activation, supporting the existence of a feed-forward loop between GSDME, caspase-3, and cytochrome c release. In contrast, the GSDME-D270A mutant failed to trigger this feedback mechanism (Fig. 6G). Together, these findings demonstrate that caspase-3-dependent GSDME cleavage is required for mitochondrial disruption and pyroptotic neuronal death in the MPTP model of Parkinson’s disease.

**Figure 6.**
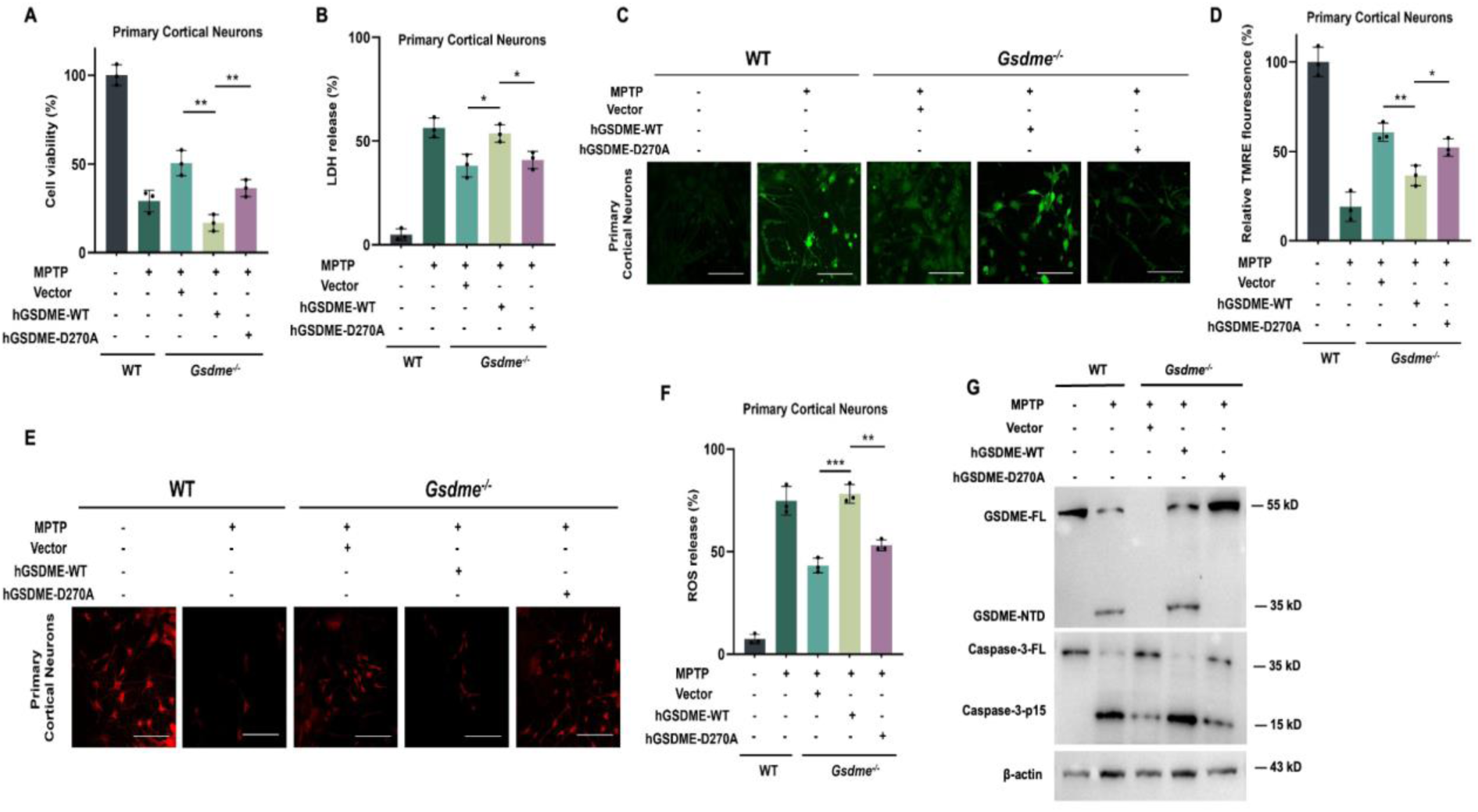
Cleavage-competent GSDME restores mitochondrial toxicity in Gsdme⁻/⁻ neurons. (**A-B**) Primary cortical neurons from WT or *Gsdme*^−/−^ mice were transduced with empty vector, WT human GSDME (hGSDME), or the cleavage-resistant D270A mutant and treated with MPTP, and assessed for cell viability (**A**) and LDH release (**B**). (**C-D**) Representative TMRE staining (**C**) and quantification of mitochondrial membrane potential (**D**) in the indicated conditions. Scale bar, 100 μm. (**E-F**) Representative DCFH-DA staining (**E**) and quantification of ROS generation (**F**) in the indicated conditions. Scale bar, 100 μm. (**G**) Immunoblots of GSDME-FL, GSDME-NTD, caspase-3-FL, caspase-3-p15, and β-actin in WT and *Gsdme*^−/−^ primary cortical neurons expressing the indicated constructs following MPTP treatment. All experiments were performed in biological triplicates. Data represent the mean ± SD. ns, not significant; *P < 0.05; **P < 0.01; ***P < 0.001; ****P < 0.0001.

## Discussion

Dopaminergic neuron loss is the defining pathology of Parkinson’s disease, yet the execution program that converts mitochondrial stress into neuron death has remained uncertain ^10,26^. Our data identify GSDME-dependent pyroptosis as a central driver of neurodegeneration in toxin-based models of Parkinson’s disease (Fig. 1-6). Across primary neurons and SH-SY5Y cells, knockdown of Gsdme or its transcriptional regulator SP1, and pharmacologic caspase-3 inhibition, blunted GSDME cleavage, preserved axons, and reduced lytic death (Fig. 1). *In vivo*, *Gsdme*-deficient mice were protected from MPTP-induced loss of dopaminergic neurons, exhibited reduced glial activation and pro-inflammatory cytokine release, and demonstrated significant improvements in motor performance, anxiety-, and depression-like behaviors (Fig. 2-3). Midbrain transcriptomics revealed that MPTP exposure induces a neuroplasticity- and development-associated transcriptional response in the substantia nigra, which was significantly attenuated in Gsdme-deficient mice, indicating that GSDME contributes to stress-induced transcriptional reprogramming likely through its role in neuroinflammation and neuronal injury amplification (Fig. 4).

Mechanistically, we show that cleaved GSDME translocates to mitochondria where it worsens organelle injury (Fig. 5-6). This localization supports a feed-forward loop in which mitochondrial damage triggers cytochrome c release and caspase-3 activation, caspase-3 generates GSDME-NT, and GSDME-NT further disrupts mitochondrial potential and raises reactive oxygen species, amplifying caspase-3 activation. Interdicting GSDME, genetically or by blocking its cleavage, breaks this loop, preserves mitochondrial structure, and limits neuronal pyroptosis. In parallel, GSDME pores at the plasma membrane produce classical pyroptotic morphology and likely augment neuroinflammation through release of intracellular contents (Fig. 1-6).

These findings integrate GSDME-mediated pyroptosis into the landscape of regulated cell death in Parkinson’s disease, which also includes BAK/BAX-dependent apoptosis and RIPK1-RIPK3-MLKL necroptosis ^11,27^. Moreover, molecular mediators of cellular rupture, such as NINJ1 ^28^, may further shape this pathogenic network. These observations support a model wherein multiple programmed death pathways operate in concert or in parallel to drive disease, underscoring the need for therapeutic strategies that can selectively intercept key nodes within this network.

The role of GSDME-dependent pyroptosis in PD also suggests broader implications for other neurodegenerative illnesses, including Alzheimer’s and Huntington’s disease. The development of GSDME-targeted interventions, such as small-molecule inhibitors of its pore-forming activity, compounds that block its caspase-3-dependent cleavage, or approaches that suppress its expression via transcriptional or RNA-based strategies, represents a promising avenue for disease-modifying therapy. Together, our findings illuminate a new path for therapeutic exploration and underscore the central role of GSDME-driven pyroptosis in neurodegenerative pathobiology.

## Methods and materials

For detailed experimental procedures, please see supplementary materials and methods.

## Acknowledgments

This work was supported by grants from the National Natural Science Foundation of China (82572011, 32161160323) and the Shanghai Committee of Science and Technology (24490713600).

## Author Contribution

J.L. conceived and designed the study. J.P., L.G., X.L., Y.W., N.F., F.X., Y.S., Y.D., Y.H., S.L., W.X., H.H., N.S. and J.W. performed the experiments and analyzed the data. J.P. and J.L. analyzed the data and wrote the manuscript. All authors discussed the results and commented on the manuscript.

## Conflict of Interest

The authors declare that they have no competing interests.

## Ethics Statement

All animal experiments were performed in accordance with the NIH Guide for the Care and Use of Laboratory Animals, with the approval of the Scientific Investigation Board of School of Life Sciences, Fudan University (2020-JS-016).

## Data availability

The RNA-seq data generated in this study were deposited in the NCBI Sequence Read Archive with accession code PRJNA1353599.

## Supplementary Information

### Methods and Materials

#### Plasmid construction and RNA interference

The human GSDME coding sequence was PCR-amplified from HeLa cDNA and cloned into the pcDNA3.1-HA expression vector. The GSDME D270A point mutant was introduced by site-directed mutagenesis using a circular PCR approach. shRNAs targeting human Sp1 (5′-GGATGGTTCTGGTCAAATACA-3′), mouse Sp1 (5′-CCTTCACAACTCAAGCTATTT-3′), human GSDME (5′-GATGATGGAGTATCTGATCTT-3′), and mouse GSDME (5′-CCGTCAAGAGAACAGTTAATA-3′) were designed, synthesized, and cloned into the pLVX-shRNA2-mCherry-Puro vector (Tsingke, Shanghai, China). All constructs were verified by Sanger sequencing.

#### Mice primary cortical neurons

Primary cortical neurons were prepared from postnatal day 0-1 C57BL/6 mice. Briefly, 12-well plates were coated with 100 μg mL^-1^ pol-L-lysine overnight and rinsed with PBS prior to use. Newborn mice were euthanized, and brains were dissected and placed in ice-cold DMEM (Hyclone) supplemented with 1% penicillin-streptomycin. Under a stereomicroscope, the olfactory bulbs, cerebellum, and meninges were removed, and cortical tissue was isolated and minced into ∼0.5-1 mm pieces. The tissue was digested with 0.25% trypsin and DNase I (Thermo Fisher) at 37 °C for up to 20 minutes, with gentle agitation every 3 minutes. Digestion was stopped by adding DMEM containing 10% FBS (Gibco), followed by centrifugation and resuspension. The dissociated cells were passed through a 70 μm strainer, counted, and plated at a density of 4-8 × 10^5^ cells/mL. After 3-4 h, non-adherent debris was removed by washing with pre-warmed serum-free DMEM, and neurons were cultured in Neurobasal medium (Thermo Fisher). At day 3, half-medium changes were performed, and from day 6 onwards, full medium changes were performed prior to experimental treatments.

#### Cell culture and treatments

Primary cortical neurons were cultured in Neurobasal medium supplemented with B-27, N-2, 0.5 mM L-glutamine, penicillin, and streptomycin (Thermo Fisher). SH-SY5Y human neuroblastoma cells were maintained in high-glucose DMEM (Hyclone) supplemented with 10% (v/v) FBS (Gibco) and penicillin-streptomycin. All cultures were kept at 37 °C in a humidified 5% CO_2_ atmosphere. For neurotoxin treatments, cells were plated in 96-well plates and exposed to 50 μM 6-hydroxydopamine (6-OHDA) and 2.5 ng mL^−1^ TNF-α (MCE), or 1 mg mL^−1^ MPTP (MCE) for 24 h. To assess caspase-3 involvement, cultures were pre-treated with the caspase-3 inhibitor Z-DEVD-FMK (MCE) at the indicated concentrations prior to neurotoxin exposure.

#### Lentiviral expression and transduction

Lentiviral particles were generated by co-transfecting HEK293T cells with the pLVX-shRNA2-mCherry-Puro or pLVX-IRES-mCherry overexpression plasmids, along with psPAX2 and pVSVG packaging plasmids, at a ratio of 8 μg:6 μg:2 μg per 10 cm dish. Viral supernatants were collected at 48 h and 72 h post-transfection, concentrated by ultracentrifugation at 30,000×g for 90 minutes, and resuspended in 1% of the original DMEM volume. For knockdown or overexpression, SH-SY5Y cells or primary neurons were cultured to ∼80% confluence and infected with concentrated virus (1% of the culture volume). After 48 h, protein expression was confirmed by immunoblotting.

#### Cell viability and cytotoxicity assays

Cell viability following neurotoxin treatments was assessed using the Cell Counting Kit-8 (CCK-8, APExBIO) according to the manufacturer’s instructions. After 24 h of treatment, the culture medium was replaced with medium containing 5% CCK-8 solution and incubated for 1 h at 37 °C in the dark. Absorbance was measured using a SpectraMax M5 plate reader. Cytotoxicity was quantified using the LDH Cytotoxicity Assay Kit (Beyotime). After treatments, supernatants were collected and incubated with LDH detection reagents (INT, diaphorase, and lactate) for 30 minutes in the dark, and absorbance was read on the SpectraMax M5.

#### Mitochondrial damage assessment

Mitochondrial membrane potential was determined after neurotoxins stimulation using the Mitochondrial Membrane Potential Assay Kit with TMRE (Beyotime) according to the manufacturer’s instructions. Briefly, after corresponding reagents treatment for 24 h, cells were incubated with 100 nM TMRE working solution at 37°C for 15∼45 min, washed with pre-warmed medium. ROS release was measured using ROS Assay Kit (Beyotime), according to the manufacturer’s instructions. Briefly, after corresponding reagents treatment for 24 h, cells were incubated with 10 μM DCFH-DA at 37 °C for 20 min, then washed with PBS. Images were captured with laser scanning confocal microscope (Olympus), and absorbance was measured on the SpectraMax M5 plate reader.

#### Protein preparation and western blot

Cells were collected, rinsed with PBS, and lysed in ice-cold RIPA buffer (50 mM Tris-HCl, pH 7.5, 150 mM NaCl, 1% Triton X-100, 1 mM EDTA) containing protease inhibitor cocktail (Roche). Lysates were centrifuged at 17,000×g for 10 minutes at 4°C, and supernatants were diluted with 5× loading buffer and separated by SDS-PAGE. Gels were transferred to nitrocellulose membrane, blocked with 5% FBS Buffer for 1 h, followed by incubation with specific antibodies: anti-Sp1 (Abcam, ab231778), anti-GSDME (Abcam, ab215191), anti-Caspase-3 (Cell Signaling Technology, 14220), anti-Tyrosine Hydroxylase (Abcam, ab137869), anti-p-α-synuclein (Abcam, ab168381), anti-Cytochrome c (Cell Signaling Technology, 4272), anti-TOM20 (Proteintech, 66777-1-Ig) and anti-β-actin (Proteintech, 66009-1-Ig). Following antibody incubation, membranes were washed with TBST and incubated with HRP labeled secondary antibody for 1h: anti-mouse IgG (Proteintech, SA00001-1) and anti-rabbit IgG (Proteintech, SA00001-2). Membranes were washed by TBST and imaged on chemiluminescent imaging system (Tanon).

#### Immunofluorescence

Primary neurons and SH-SY5Y cells were seeded onto 35 mm confocal dishes and subjected to transfection or drug treatment as required. Cells were fixed with 4% paraformaldehyde for 30 min at room temperature and washed three times with PBS. After fixing, cells were incubated with primary antibodies diluted in blocking buffer containing FBS, TritonX-100 and anti-Tuj1 antibody (Abcam, ab52653), anti-GSDME antibody (Proteintech, 13075-1-AP) or TOMM20 (Abcam, ab186735) overnight at 4 °C. Cells were washed with PBST and incubated with secondary antibodies (Beyotime A0468, Abcam ab150113, Abcam ab150078) diluted in PBST for 1 h in the dark. After extensive washing with PBST and PBS, cells were counterstained with DAPI mounting medium, covered with a coverslip, and stored at 4 °C in the dark. Fluorescence images were acquired using a laser scanning confocal microscope (Olympus).

#### MPTP-induced Parkinson’s disease model and behavioral tests

##### Establishing the MPTP-induced PD model

Around 8-week-old WT and *Gsdme*^⁻/⁻^ mice were divided into control and experimental groups (n=10∼15). Experimental mice received intraperitoneal injections of MPTP (20 mL/kg, 1 g/L in saline) every 2 h for four doses, while controls received saline. Behavioral tests were performed one week after the final injection.

##### Open field test

Mice were individually placed in an open-field chamber and allowed to explore freely for 5 min. The video tracking system Noldus EthoVision was used to record locomotor trajectories, total distance traveled, and entries/time spent in the central zone. Chambers were cleaned with 75% ethanol between trials. All behavioral data were exported for statistical analysis.

##### Rotarod test

Mice were trained for three consecutive days using a rotating rod apparatus. On day 1, they were placed on the rod at a constant speed of 4 rpm for 5 min; mice that fell were gently placed back on the rod. On days 2 and 3, the rotation accelerated from 4 to 40 rpm for 5 min. During the test, the rod accelerated from 4 to 40 rpm, and latency to fall was recorded without repositioning fallen mice. The apparatus was cleaned thoroughly with distilled water between trials.

##### Pole test

Mice were placed head-up on top of a vertical gauze-wrapped rod. After turning, they descended to the bottom. Mice were trained for two days, then tested on the third day. Time taken to descend was recorded. The apparatus was cleaned between trials.

##### Tail suspension test

Mice were suspended by the tail using adhesive tape fixed to a clamp, with heads hanging freely. Struggling behavior was recorded for 5 minutes, and the total duration of immobility and active movement was analyzed.

##### Elevated plus maze test

Mice were individually placed in an elevated plus maze and allowed to explore freely for 8 min. The video tracking system Noldus EthoVision was used to record locomotor trajectories, total distance traveled, and entries/time spent in the open and closed arm. Maze was cleaned with 75% ethanol between trials. All behavioral data were exported for statistical analysis.

##### Y maze test

Mice were individually placed in a Y maze and allowed to explore freely for 10 min. The video tracking system Noldus EthoVision was used to record locomotor trajectories and spontaneous alternation among the three arms. Maze was cleaned with 75% ethanol between trials. All behavioral data were exported for statistical analysis.

#### Immunohistochemistry

Mice were euthanized by cervical dislocation, and brains were carefully extracted after removing the scalp and skull. The cortex, substantia nigra, and striatum were dissected, fixed in 4% paraformaldehyde, and submitted for paraffin embedding, sectioning, and mounting (Servicebio). Sections were deparaffinized in xylene for 20 min, rehydrated through graded ethanol (100%, 95%, 90%, 85%, 80%, 75%, 10 min each), and rinsed in tap water. Antigen retrieval was performed using 0.01 M citrate buffer in a microwave oven, with two cycles of 30 s boiling and natural cooling. Endogenous peroxidase activity was blocked using 3% hydrogen peroxide for 20 min, followed by PBS washes. Sections were blocked with 10% goat serum for 1 h, incubated overnight at 4 °C with primary antibodies: anti-Tyrosine Hydroxylase (Abcam, ab137869), anti-GSDME (Abcam, ab215191), anti-GFAP (Abcam, ab4674), anti-CD11b (Abcam, ab133357) and Iba1 (Abcam, ab5076). After washed with PBS and incubated with secondary antibodies at 37 °C for 1 h and further PBS washes, sections were treated with SABC reagent for 30 min. Visualization was done with DAB for 5 min, followed by hematoxylin counterstaining for 2 min. Slides were dehydrated through graded ethanol and cleared in xylene. Finally, coverslips were mounted with neutral resin, and images were acquired using a light microscope.

#### RNA sequencing and data analysis

Total RNA was extracted from mouse brain tissues of WT, WT-MPTP and *Gsdme*^⁻/⁻^-MPTP mice using TRIzol reagent. Libraries were prepared using a standard Illumina protocol and sequenced on the NovaSeq 6000 platform. Clean reads were aligned to the mouse reference genome (GRCm39) using HISAT2. Differential expression analysis was performed with DESeq2, with significantly changed genes defined as those with |log₂FC| ≥ 1 and adjusted p < 0.05. GO and KEGG enrichment analyses were conducted using ClusterProfiler in R.

#### Quantitative real-time PCR (qPCR)

Total RNA was extracted from mouse substantia nigra tissues using TRIzol reagent, and 1 μg of RNA was reverse-transcribed into cDNA using a HiScript III RT SuperMix Kit (Vazyme). Quantitative PCR was performed using ChamQ SYBR qPCR Master Mix (Vazyme) on a QuantStudio 6 Flex Real-Time PCR System (Applied Biosystems). Gene-specific primers were designed using Primer-BLAST, and β-actin was used as an internal control. Relative gene expression was calculated using the method.

### Statistical analysis

All experiments were independently repeated at least three times. Data were analyzed using GraphPad Prism 9.0 (GraphPad Software Inc., USA) and are expressed as mean ± SD. Statistical significance was evaluated using Student’s t-test, one-way ANOVA, or two-way ANOVA, as appropriate. A p-value < 0.05 was considered statistically significant.

---

**Supplementary Figure 1.**
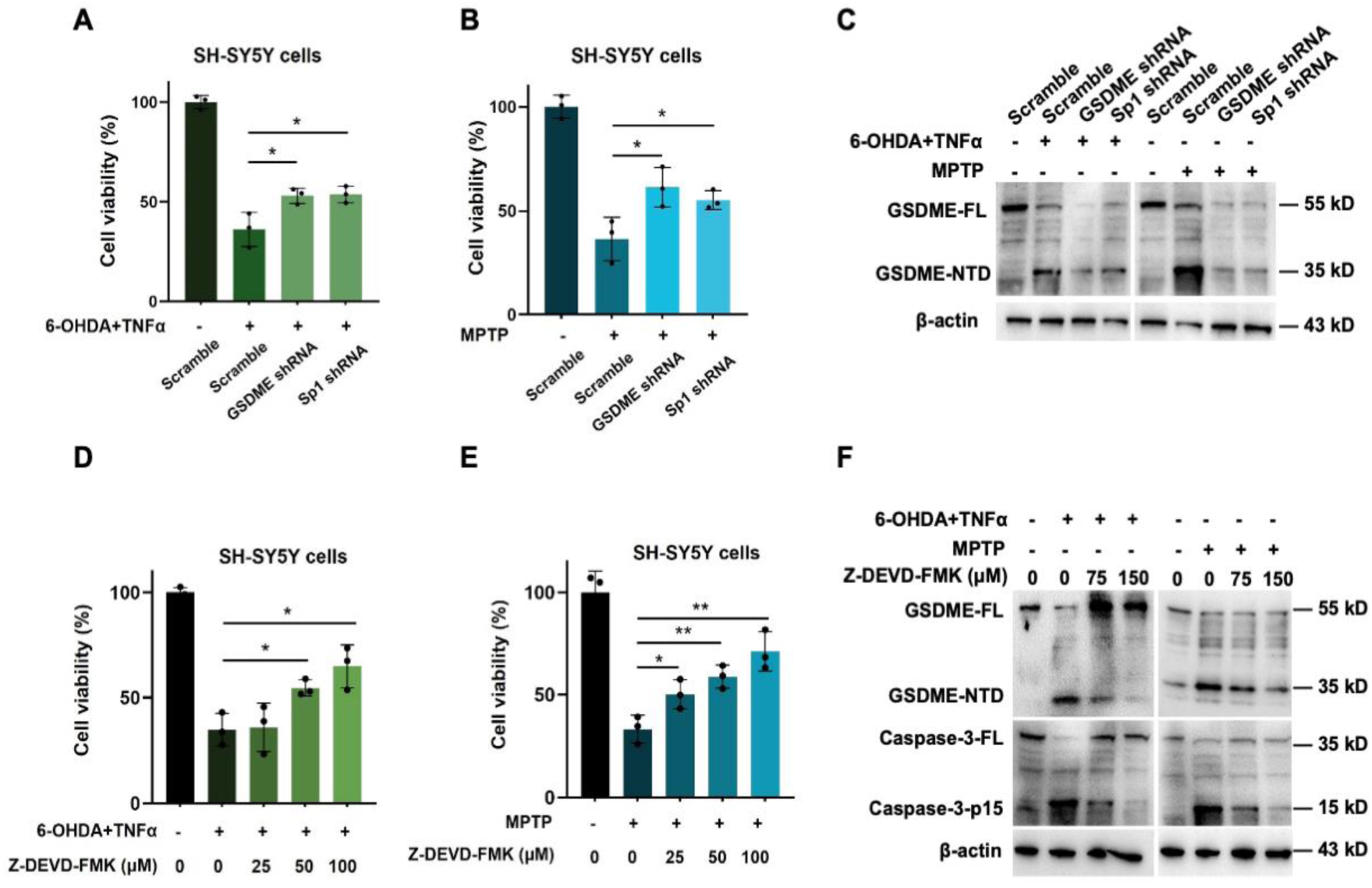
Genetic or pharmacological inhibition of GSDME reduces neurotoxin-triggered pyroptosis. (**A-C**) Human SH-SY5Y cells were transduced with lentivirus encoding scrambled control, GSDME shRNA, or Sp1 shRNA. After 72 h, cultures were challenged with 6-OHDA (+ TNF-α, 10 ng ml⁻¹) or MPTP as indicated. Cell viability was quantified 24 h later (normalized to vehicle) (**A, B**). Immunoblots were performed for full-length GSDME (GSDME-FL), cleaved N-terminal GSDME (GSDME-NTD), and β-actin (**C**). (**D-F**) Neurotoxin-treated SH-SY5Y cells were exposed to increasing concentrations of the caspase-3 inhibitor Z-DEVD-FMK, and cell viability was quantified (**D, E**). Immunoblot analyses of GSDME-FL, GSDME-NTD, caspase-3-FL, and caspase-3-p15 (with β-actin as a loading control) in Z-DEVD-FMK–treated, neurotoxin-stimulated cells (**F**). All experiments were performed in biological triplicates. Data represent the mean ± SD. ns, not significant; *P < 0.05; **P < 0.01; ***P < 0.001; ****P < 0.0001.

## Notes

### Competing Interest Statement

The authors have declared no competing interest.

